# Loss of *Frrs1l* disrupts synaptic AMPA receptor function, and results in neurodevelopmental, motor, cognitive and electrographical abnormalities

**DOI:** 10.1101/388561

**Authors:** Michelle Stewart, Petrina Lau, Gareth Banks, Rasneer Sonia Bains, Enrico Castroflorio, Peter L. Oliver, Christine L. Dixon, Michael C. Kruer, Dimitri M. Kullmann, Abraham Acevedo-Arozena, Sara E. Wells, Silvia Corrochano, Patrick M. Nolan

**Affiliations:** MRC Harwell Institute, Harwell Campus, Oxfordshire OX11 0RD, UK; University College London (UCL) London, UK; Barrow Neurological Institute, Phoenix Children’s Hospital, Phoenix, USA; Unidad de Investigación, Hospital Universitario de Canarias, Fundación Canaria de Investigación Sanitaria and Instituto de Tecnologías Biomédicas (ITB), Tenerife, Spain.

## Abstract

**Summary statement:** In this study, we show that the loss of the epilepsy-related gene *Frrs1l* in mice causes a dramatic reduction in AMPA receptor levels at the synapse. This change elicits severe motor and coordination disabilities, hyperactivity, cognitive defects, behavioural seizures and abnormal electroencephalographic (EEG) patterns.

**Abstract:** Loss of function mutations in the human AMPA receptor-associated protein, ferric chelate reductase 1-like (FRRS1L), are associated with a devastating neurological condition incorporating choreoathetosis, cognitive deficits and epileptic encephalopathies. Furthermore, evidence from overexpression and *ex vivo* studies have implicated FRRS1L in AMPA receptor biogenesis and assembly, suggesting that changes in glutamatergic signalling might underlie the disorder. Here, we investigated the neurological and neurobehavioural correlates of the disorder using a mouse *Frrs1l* null mutant. The study revealed several neurological defects that mirrored those seen in human patients. We established that mice lacking *Frrs1l* suffered from a broad spectrum of early-onset motor deficits with no progressive, age-related deterioration. Moreover, *Frrs1l^-/-^* mice were hyperactive irrespective of test environment, exhibited working memory deficits and displayed significant sleep fragmentation. Longitudinal electroencephalographic recordings also revealed abnormal EEG in *Frrs1l^-/-^* mice. Parallel investigations into disease aetiology identified a specific deficiency in AMPA receptor levels in the brain of *Frrs1l^-/-^* mice, while the general levels of several other synaptic components remained unchanged with no obvious alterations in the number of synapses. Furthermore, we established that *Frrsl1* deletion results in glycosylation deficits in GLUA2 and GLUA4 AMPA receptor proteins, leading to cytoplasmic retention and a reduction of those specific AMPA receptor levels in the postsynaptic membrane. Overall, this study determines, for the first time *in vivo*, how loss of FRRS1L function can affect glutamatergic signalling and provides mechanistic insight into the development and progression of a human hyperkinetic disorder.

## Introduction

Ferric chelate reductase 1 like (FRRS1L) is a novel, highly conserved, brain specific protein whose functional characterisation has only recently been under investigation. Studies in patients have found nine families with recessive mutations in *FRRS1L* which result in severe intellectual disability, movement disorders, hypotonia and epilepsy (Madeo *et al.*, 2016; Shaheen *et al.*, 2016; Brechet *et al.*, 2017). In some patients, these clinical symptoms are accompanied by neurodegeneration in the cortex and cerebellum. Several families have now been diagnosed with this devastating condition, arguing for the inclusion of this gene in the diagnostic screening for epilepsy and dyskinetic disorders (Carecchio and Mencacci, 2017). The dramatic clinical consequences of carrying mutations in this gene point to an important neurological function for *FRRS1L*, which has not yet been elucidated, hence challenging efforts in therapeutic development.

Although named for its sequence similarity to ferric chelate reductase 1, FRRS1L has only a poorly characterised DOMON domain (DOpamine beta-MOnooxygenase N-terminal domain, IPR005018) and a transmembrane domain, with the ferric chelate reductase domain being absent, therefore its function is likely to be distinct from that of its namesake, FRRS1. *Frrs1l* is expressed in the central nervous system (CNS) and testis of adult mice and in developing embryonic forebrain (Madeo *et al.*, 2016). Further expression analysis in the adult mouse brain, shows *Frrs1l* expression in the excitatory neurons in the cerebral cortex, hippocampus and midbrain, medium spiny neurons in the striatum, granule cells in the dentate gyrus and Purkinje cells in the cerebellum (Zeisel, A *et al.,* 2018 preprint). Emerging studies have begun to unravel the role of FRRS1L in the CNS, importantly in AMPA receptor complex function. FRRS1L co-localises with Calnexin in the endoplasmic reticulum (ER) of rat hippocampal neurons (Brechet *et al.*, 2017). Results of knockdown and exogenous overexpression studies in cultured hippocampal neurons suggest that FRRS1L, along with carnitine palmytoyltransferase 1c (CPT1C), is involved in the early stages of AMPA receptor complex biogenesis, binding to the core AMPA proteins, GLUA1-4, but dissociating before the final auxiliary proteins bind to make a functional receptor (Brechet *et al.*, 2017). Furthermore, reduction of FRRS1L levels in cultured hippocampal neurons leads to an overall decrease in AMPA receptor levels, as well as to modifications in synaptic transmission. In addition, interactions with dynein complex proteins suggest a potential role for FRRS1L in dynein based AMPA trafficking (Han *et al.*, 2017).

AMPA receptors are essential ionotropic glutamate receptors and mediate much of the fast-excitatory synaptic transmission in the brain. AMPA receptors are composed of four core proteins, GLUA1-4, which form a heterotetrameric complex at the centre of the receptor. Associated with this core complex are a variety of auxiliary subunits with distinct roles in the maturation of AMPA receptors (Chen *et al.*, 2000; Tomita *et al.*, 2003; Kato *et al.*, 2010; Schwenk *et al.*, 2012, 2014; Erlenhardt *et al.*, 2016). These auxiliary proteins have distinct roles in regulating the spatio-temporal activity of AMPAs, however many of these roles have yet to be elucidated.

The majority of human variants in patients with homozygous mutations in *FRRS1L* are predicted to lead to a premature stop codon and loss of the transmembrane domain, consequently leading to a loss of function. A knockout of the murine *Frrs1l* gene (*Frrs1l^tm1b/tm1b^*) has been generated by the International Mouse Phenotyping Consortium (IMPC, http://www.mousephenotype.org/) to investigate the consequences of *Frrs1l* loss *in vivo*. Initial characterisation of this line uncovered a range of aberrant phenotypes including hyperactivity, abnormal gait, decreased grip strength and partial pre-weaning lethality (Koscielny *et al.*, 2014; *mousephenotype.org*, accessed 01-06-2018). In the current study, we make use of this mouse line to explore its validity as a pre-clinical model and to investigate the phenotypic deficits in more depth. Moreover, we make use of this model to study whether a disturbance in AMPA receptor maturation is the mechanism underlying the pathology of the disorder providing new *in vivo* evidence for the pivotal role that *Frrs1l has* in AMPA receptor physiology.

## Results

### Loss of Frrs1l results in increased neonatal lethality, smaller size and early onset motor deficits

*Frrs1l*^-/-^ mice are born at expected Mendelian ratios; milk is present in the stomach, breathing is apparently normal and it is not possible to visibly distinguish between *Frrs1l*^-/-^ and wild-type littermates. However, >90% of *Frrs1l*^-/-^ neonates die between 12 and 24 hours after birth. Analysis of numbers per genotype at weaning show a difference in expected ratios (*p<0.0001*), whereas at postnatal day 0 (P0) the ratio of genotypes is not significantly different to that expected (*p=0.43*) (Fig. 1A). Tissue was collected from any pups that were found dead in the first days after birth and genotyping carried out. We found mortality to be higher in *Frrs1l*^-/-^ with a greater proportion of homozygotes being found dead than would be expected by chance if this was not a genotype effect (*p<0.001*). Gross pathology was performed on pups at P0 and no obvious abnormalities were found in 44 tissues examined (data not shown). *Frrs1l*^-/-^ mice that survive past P2 continue to thrive to weaning and beyond. Five of the nine female homozygous mice were killed during the course of the study as they reached previously specified humane endpoints, including; seizure without full recovery (n=1), self-inflicted wounds and stereotypical behaviour (n=2), uncoordinated gait impinging on the ability to feed (n=1) and breathing difficulties (n=1).

**Fig. 1.**
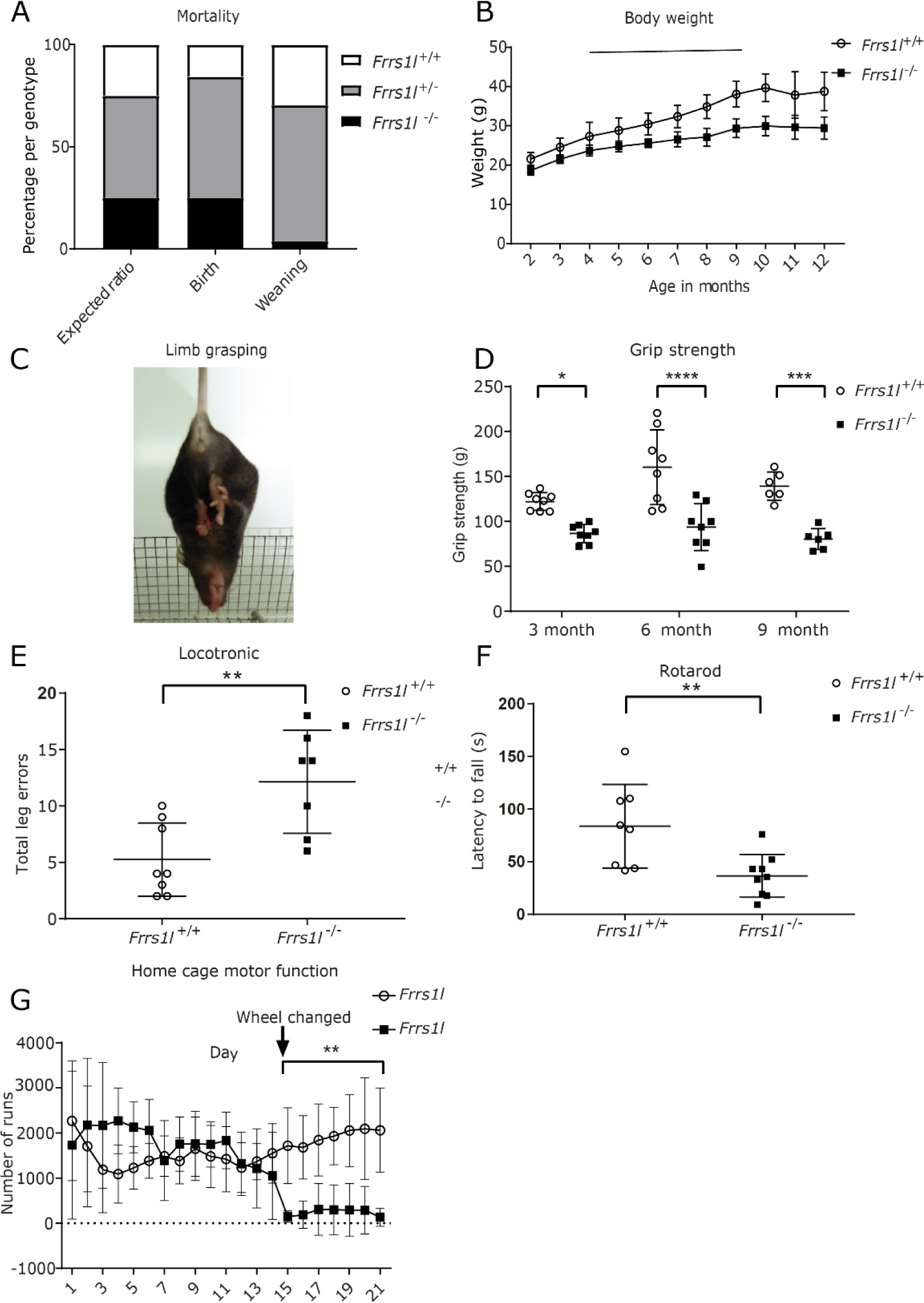
Frrs1l^-/-^ have reduced survival, decreased body weight, coordination and limb grasping abnormalities. Frrs1l^-/-^ are born in accordance with predicted mendelian ratios (P=0.43) and show no significant difference to expected numbers. Pups were genotyped at weaning and greatly reduced numbers of Frrs1l^-/-^ were found (p<0.0001 (A)). Data analysed by chi-square test. Frrs1l^-/-^ have significantly lower weight from 6 months of age but show a similar weight curve and no deterioration (B). Data analysed by two way ANOVA with post-hoc comparisons (*p<0.05, **p<0.01, ***p<0.001). n=7 Frrs1l^+/+^ from 2-11 months and n=5 Frrs1l^+/+^ at 12 months, n=9 Frrs1l^-/-^ at 2-4 months, n=8 Frrs1l^-/-^at 5 months, n=7 Frrs1l^-/-^ at 6-8 months, n=6 Frrs1l^-/-^ at 9-12 months. Frrs1l^-/-^ show increased incidence of limb grasping (p<0.01 at 3, 6 and 9 months), representative image (C). Grip strength of Frrs1l ^-/-^ is significantly reduced at three, six and nine months when compared to wild-type littermate controls **(D)**. Data analysed by repeated measures ANOVA followed by Sidak multiple comparisons test, data show is average of three trials at each time point. n=8 Frrs1l ^+/^; n=8 Frrs1l ^-/-^ at 3 and 6 months, n=6 Frrs1l^+/+^; n=6 Frrs1l^-/-^ at 9 months. Misplacement of feet on the Locotronic horizontal ladder results in an increased number of errors in Frrs1l^-/-^ **(E)**, n=8 Frrs1l ^+/+^; n=7 Frrs1l ^-/-^ analysed using general linear model with Poisson distribution. Frrs1l^-/-^ have a significantly decreased latency to fall from an acceleration rotarod (p<0.01) (n=8 Frrs1l ^+/+^; n=8 Frrs1l ^-/-^). Data analysed by repeated measures ANOVA. Frrs1l^-/-^ display no difference in wheel running on simple wheels but show a significant difference when changed to complex wheels, analysed using ANOVA followed by Sidak multiple comparisons test n=5 Frrs1l ^+/+^; n=5 Frrs1l ^-/-^**(G)**.(*p<0.05, **p<0.01, ***p<0.001). Body weight (B) and Wheel running (G) data are mean and standard deviation.

In order to confirm that homozygous mutant mice no longer express *Frrs1l*, we conducted qPCR assays with primers spanning all five exons of the gene, including the targeted exon 3. We confirmed no significant expression of *Frrs1l* in P0 and adult *Frrs1l*^-/-^ brains compared to wild-type littermates (Fig. S1A and B). Of the *Frrs1l^-/-^* mice that survive to weaning, all show prominent reduced body weight when compared to littermate controls (*P<0.05* from 6 months onwards) (Fig. 1B). However, the weight curve is not dissimilar to wild-type and does not show significant decline with age up to 14 months, suggesting a neurodevelopmental effect rather than a progressive wasting phenotype. Given the difference in body size, we further examined archived IMPC x-ray images and found that *Frrs1^-/-^* mice have a significantly shorter tibia length, and therefore smaller body size, than wild-type controls in both females (P<0.01) and males (P<0.05) (http://www.mousephenotype.org, 2015). IMPC data also show no differences in body composition or calorimetric measurements of metabolic rate. In some cases, several years after onset of symptoms, human patients carrying mutations in *FRRS1L* express cerebellar atrophy and other pathological alterations in the brain (Madeo *et al.*, 2016). In *Frrs1l*^-/-^ cohorts, we found total brain weight to be approximately 10% less than wild-type controls (*Frrs1l^+/+^* 0.459 ± 0.018 and *Frrs1l*^-/-^ 0.417 ± 0.009) (p<0.05), however no gross anatomical pathologies were evident Moreover, brain size differences were not significant after normalisation for body weight.

Animals that survived through early postnatal development were assessed using a focused battery of physiological, behavioural and motor function tests throughout adulthood. Motor phenotyping was carried out in order to determine whether *Frrs1l*^-/-^ mice expressed abnormalities in movement and muscle force similar to those described in humans carrying *FRRS1L* mutations. We used a standard battery of tests including SHIRPA, grip strength and rotarod, followed by more complex testing of motor function using a horizontal ladder and a three week trial that measures progressive wheel running performance parameters. On visual inspection, *Frrs1l*^-/-^ mice have an abnormal gait which is evident at weaning and, in SHIRPA testing, show additional significant differences from wild-type littermate controls when assessed at 3, 6 and 9 months of age (*P<0.05, P<0.01 and P<0.001* respectively) (Table 1). Abnormalities highlighted in the SHIRPA test include; lack of co-ordination, demonstrated by inability to place feet correctly on a grid floor (*p<0.001* at 9 months), loss of grip when climbing down a vertical grid (*p<0.001* at 9 months), and muscle weakness indicated by an increased incidence of limb grasping (*p<0.01* at all time points) (Table.1 and Fig.1C). Limb grasping is also associated with defects in many neurological disorders, and it may indicate alterations in cortico-striatal circuits (Lalonde and Strazielle, 2011).

**Table 1.**
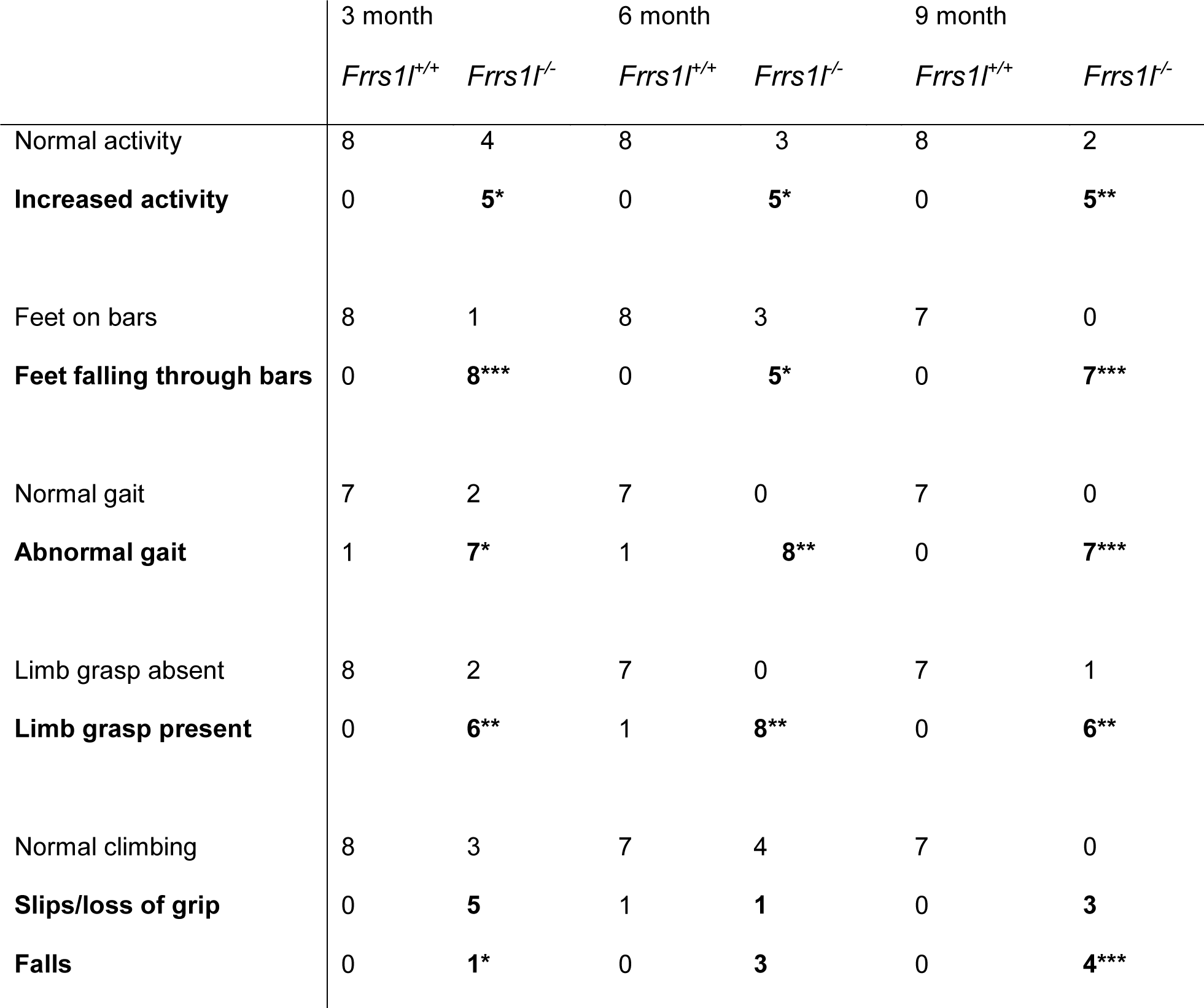
Longitudinal SHIRPA analysis shows co-ordination deficits from an early age. From 3 months of age Frrs1l^-/-^ show significantly increased activity in the viewing jar, increased incidence of feet falling through the bars, abnormal gait, limb grasping and inability to climb down a wire mesh without slipping (negative geotaxis). Deterioration up to the age of 9 months is not noticeable except that there is an increase in falling during negative geotaxis. Number of animals with each condition in relevant columns (*p<0.05, **p<0.01, ***p<0.001)

We measured grip strength at 3, 6 and 9 months of age to confirm quantitatively that the loss of *Frrs1l* resulted in muscle weakness. Indeed, grip force in all four limbs was significantly reduced in *Frrs1l*^-/-^ mice compared to wild-type controls (Fig. 1D). The effect remained significant even after correcting for weight differences (3 months *p<0.05*, 6 months *p<0.05*, 9 months *p<0.01*). This loss of muscle strength is expressed from an early age without further deterioration, suggesting a developmental non-progressive phenotype.

Next, we assessed gait and motor co-ordination by performing three complementary tests at a single time point for each: a horizontal ladder challenge (Locotronic), a rotarod test and a wheel running paradigm. In the Locotronic challenge, *Frrs1l*^-/-^ mice showed an increased frequency of errors (misplacement of feet) whilst moving along the horizontal ladder when compared to their littermate controls (*p<0.01*) (Fig. 1E). This elaborates upon the previous observations in the SHIRPA test where homozygotes had a higher number of instances of feet falling through the bars of the grid.

Supporting these data, *Frrs1l*^-/-^ mice also had a shorter latency to fall when placed on an accelerating rotarod (*p<0.05*) (Fig. 1F). In the motor function assessment by wheel running, no differences were observed between *Frrs1l*^-/-^ and wild-type mice during the first two weeks. In the third week, the standard wheel was removed and replaced by a complex wheel with rungs missing at uneven intervals. After the new challenge was introduced *Frrs1l*^-/-^ were unable to run at all, showing a drop in running attempts to almost zero for the remainder of the third week (*p<0.001*) (Fig. 1G).

Thus, all tests evidence that mice lacking the *Frrs1l* gene suffer from a dramatic loss in muscle strength with loss of motor co-ordination and motor disabilities from an early age.

### Frrs1l^-/-^ mice are hyperactive, have cognitive deficits and abnormalities in sleep

Previous IMPC-based assessment of these mice had indicated a hyperactivity phenotype. To confirm and further define this hyperactivity, mice were assessed at a number of time-points in group-housed conditions in the home cage to evaluate their activity continuously in an undisturbed, non-stressful environment. *Frrs1l*^-/-^ mice displayed increased activity when recorded at 10 weeks of age throughout both light and dark phases (*p<0.05* light, *p<0.0001* dark) and increased activity at six and nine months in the dark phase only (*p<0.05* at both time points) (Fig. 2A and B). These home cage data indicate the hyperactivity is not a consequence of being introduced to a novel environment.

**Fig. 2.**
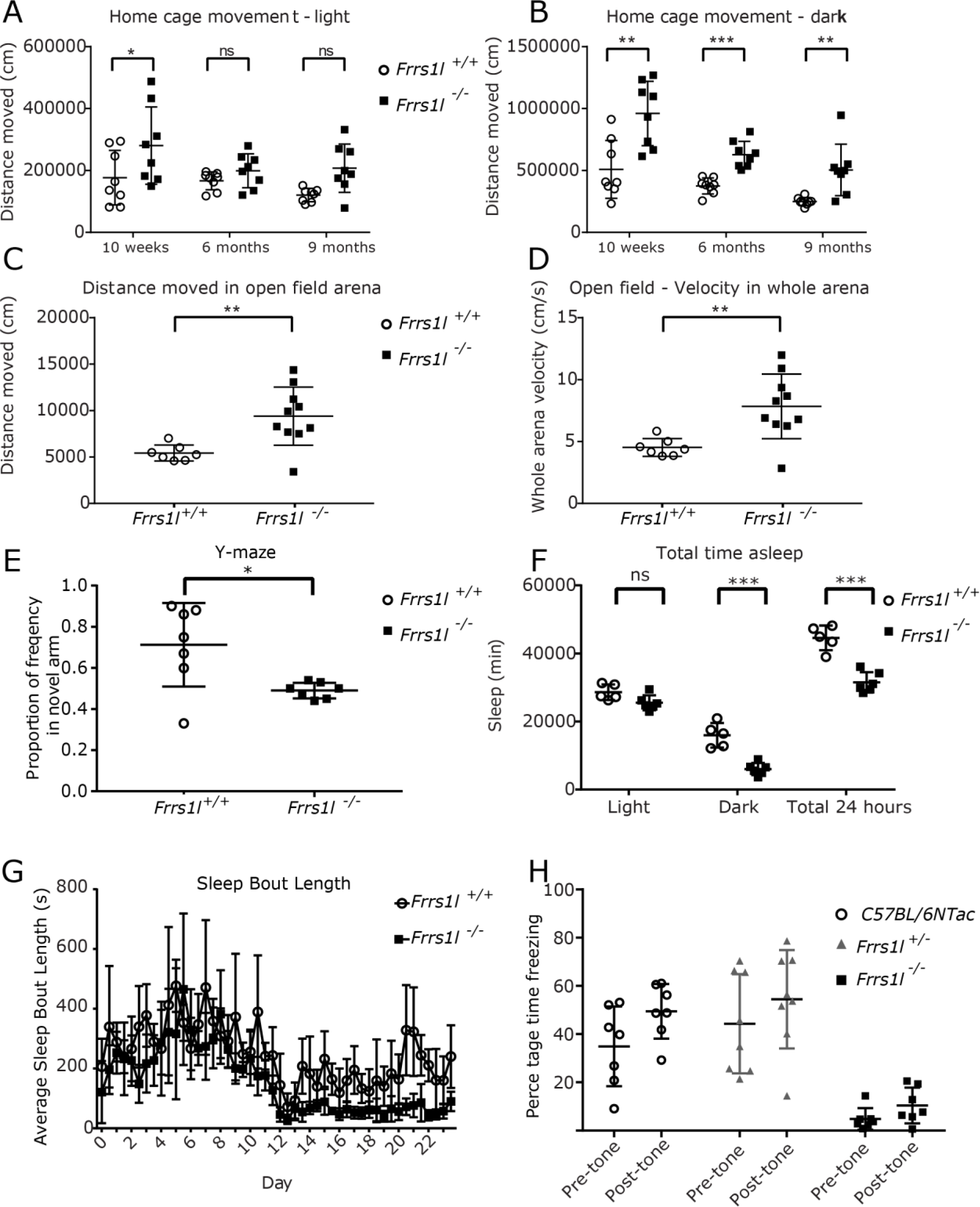
Deletion of Frrs1l causes hyperactivity, working memory deficits and abnormal sleep pattern. An increase in distance moved is apparent in the home cage both during the light phase and the dark phase at 10 weeks old (p<0.05), and in the dark phase only at 6 months and 9 months (p<0.01) (A and B). Data analysed using ANOVA followed by Sidak multiple comparisons test, n=8 Frrs1l^+/+^, n=8 Frrs1l^-/-^. Total distance moved (C) and velocity in an open field arena (D) are significantly increased in Frrs1l^-/-^ when compared to Frrs1l^+/+^ controls (p<0.01). Data analysed by t-test. n=8 Frrs1l^+/+^; n=8 Frrs1l^-/-^. (E) Frrs1l^-/-^ do not show a preference for the novel arm in a forced alternation y-maze task. Frequency of Frrs1l^-/-^ entry into the novel arm stays at chance level, while Frrs1l^+/+^ show increased exploration. Data are ratio of frequency in novel arm (frequency in novel arm/(frequency in novel arm-frequency in familiar arm) n=7 Frrs1l^+/+^; n=7 Frrs1l^-/-^.(*p<0.05). Data analysed using t-test. (F) Frrs1l^-/-^ exhibit abnormal sleep behaviour. Frrs1l^-/-^ show decreased total time asleep in a 24 hour period, with the total time asleep in the dark phase being significantly reduced (p<0.001), Sleep bout length is significantly reduced (G). Demonstrating that Frrs1l^-/-^ have a fragmented sleep pattern, with less time asleep overall and shorter sleep bouts Frrs1l^+/+^; n=5, Frrs1l^-/-^ n=6 (*p<0.05, **p<0.01, ***p<0.001). For average sleep bout length (G) data are mean and standard deviation. A separate cohort of Frrs1l^-/-^ males, assessed as part of the IMPC pipeline, do not show an increase in freezing in response to tone during a cued fear conditioning paradigm. (H) Difference in pre-cue and post cue freezing was significantly lower compared to controls (p=0.004).

In the novel environment of the open field test at a single time point, *Frrs1l*^-/-^ mice exhibited increased activity in the whole arena demonstrated by a greater total distance moved (*p<0.01*) (Fig. 2C) and an increased velocity (*p<0.01*) (Fig. 2D), supporting the hyperactivity phenotype observed in the home cage analysis. The frequency to enter the centre of the arena and velocity in the centre of the arena were also significantly increased (*p<0.05*), however distance moved and duration in the centre were not different between *Frrs1l*^-/-^ and wild-type, indicating that the mice are hyperactive but that this is not necessarily related to an altered anxiety state in mutants (Fig. S2A).

Learning disabilities are among the common features associated with intellectual disability, such as those seen in patients carrying mutations in *FRRS1L*. In the null mice, we evaluated working memory using the Y-maze forced alternation test. Interestingly, *Frrs1l*^-/-^ mice showed no preference for the novel arm (Fig. 2E) (*p<0.05*) suggesting a working memory deficit. Previous studies have demonstrated that defects in AMPA receptor composition are associated with both intellectual disability and perturbed sleep patterns (Davies *et al.*, 2017). Since FRSS1L has been proposed to play a role in AMPA receptor assembly, we assessed sleep status (immobility-defined sleep) in *Frrs1l*^-/-^ animals using passive infrared movement tracking (PIR)(Brown *et al.*, 2016). The total sleep of *Frrs1l*^-/-^ animals was significantly less than wild-type controls in the dark phase of the light cycle (time spent asleep in dark *p=0.0002*) (Fig. 2F). Additionally, the average length of sleep bouts was significantly reduced in *Frrs1l*^-/-^ animals (average sleep bout length p=0.001). Analysis of sleep bout length in the light and dark phases of the light cycle revealed that *Frrs1l*^-/-^ animals had a significant reduction in sleep bout length in the dark phase of the light cycle, with no significant effect in the light phase (average sleep bout length in dark p=0.00001, average sleep bout length in light p=0.079) (Fig. 2G). Interestingly, *Frrs1l*^-/-^ mice show no overt changes in circadian period or entrainment (data not shown).

We also examined the IMPC archive data on the Fear Conditioning paradigm, a test used to measure cognitive abilities, specifically those associated with non-declarative memory formation (LaBar and Cabeza, 2006). Mice lacking FRRS1L have deficits in cued but not contextual fear conditioning, demonstrating an inability to learn the association between a tone and an aversive stimulus and therefore an impairment in implicit memory (Fig 2H).

### Loss of *Frrs1l* causes abnormal EEG

Mutations in *FRRS1L* are associated with epileptic encephalopathy. One mouse died during a generalized convulsion. We observed episodes of behavioural arrest and lordotic posture in other *Frrs1l*^-/-^ animals. Therefore we carried out EEG recordings in *Frrs1l*^-/-^ animals to ascertain whether these episodes represent seizures. After the implantation of EEG transmitters in adult mice (n=2 *Frrs1l^/+/+^*, n=3 *Frrs1l^-/-^,* and n=2 C57BL/6NTac), we recorded their EEG activity in their home-cage environment for 5 to 15 days.

*Frrs1l*^-/-^ mice exhibited frequent runs of high amplitude delta activity that were not present in control animals (C57BL/6NTac and *Frrs1l^/+/+^* animals) (Fig. 3A and B). Although we did not capture discrete electrographic seizures, these results are consistent with the view that loss of *Frrsl1* leads to a profound encephalopathy.

**Fig. 3.**
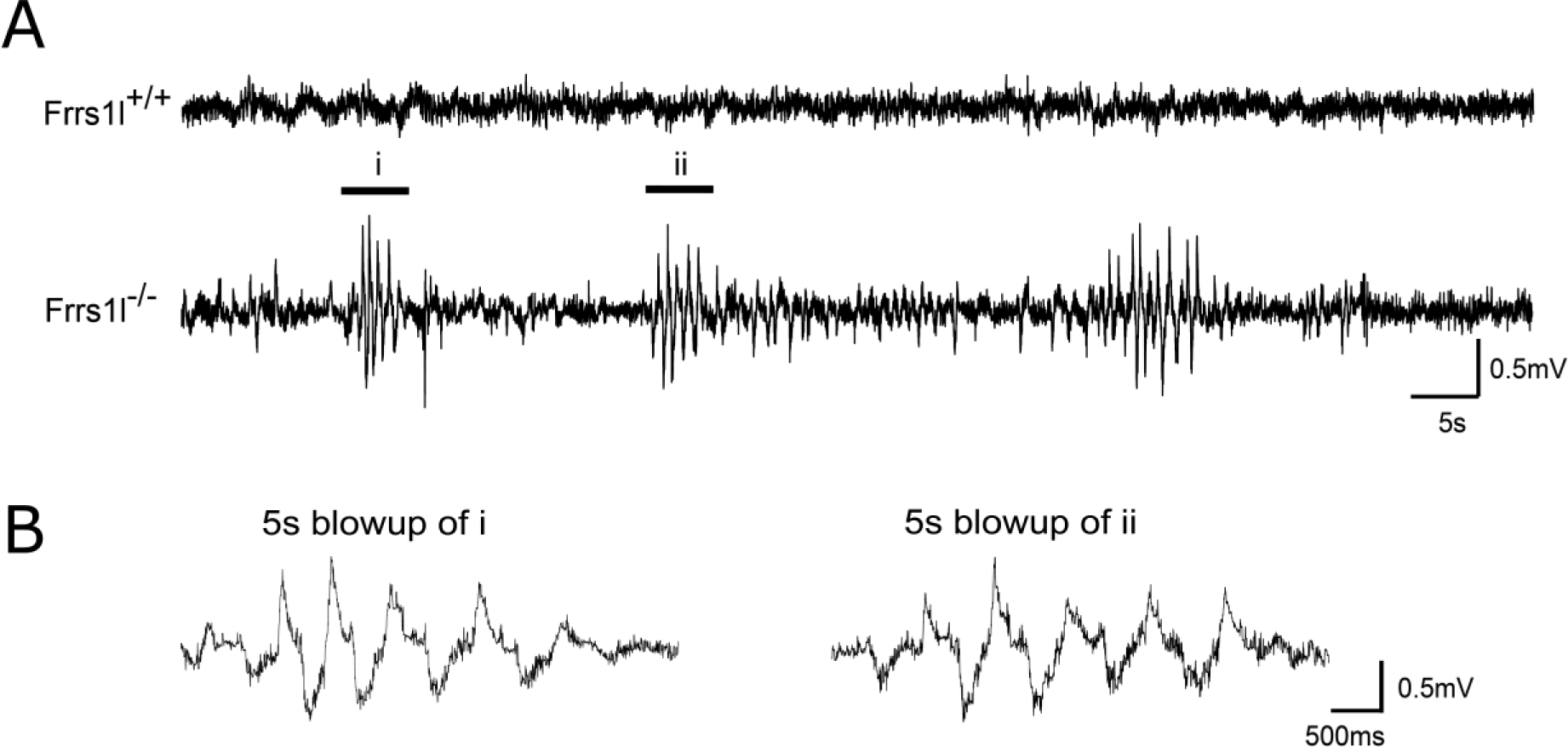
Abnormal delta frequency EEG activity in Frrs1l^-/-^animals. EEG recorded in representative Frrs1l^+/+^ and Frrs1l^-/-^ animals are shown in 100s windows. (A). Sample EEG traces were extracted and shown 100s windows. Large amplitude sawtooth-like spike trains were observed at delta frequency (ranging between 0.5 Hz – 2Hz) in two of the mutants while abnormal high theta oscillation (ranging between 6 Hz – 8Hz) was reported in the third Frrs1l^-/-^ animal. (B). Two 5s representative spike trains were extracted and enlarged from recording in the Frrs1l^-/-^ shown in (A). The waveform showed a dominant peak concentrated at 1 Hz – 2 Hz, delta frequency after Fast Fourier Transform of traces from (B), data not shown.

### Decreased AMPA receptor protein levels in Frrs1l^-/-^ brain

In order to understand the mechanisms underlying the neurological and behavioural deficits described above, we investigated whether AMPA receptor levels were altered *in vivo* as a consequence of *Frrs1l* deletion, as suggested in earlier *in vitro* studies (Brechet *et al.*, 2017)). We first examined whether the deletion of *Frss1l* would cause alterations in the gene expression levels of the four core AMPA receptor genes (*Gria1, Gria2, Gria3 and Gria4*) and found no differences between wild-type and *Frrs1l*^-/-^ mice either in P0 brain or in adult brain (Fig.S1C and Fig.4A). We next examined levels of three core AMPA receptor proteins (GLUA1, GLUA2 and GLUA4) in P0 brain and in 14 month-old cerebellum. In adult cerebellum, GLUA1, GLUA2 and GLUA4 levels were all significantly reduced in *Frrs1l*^-/-^ compared to wild-type controls (*p<0.001, p<0.01, p<0.001* respectively) (Fig. 4B and C). In P0 brain only GLUA1 was significantly reduced, while alterations in immunoreactive band mobility were noted for both GLUA2 and GLUA4 (Fig. S3B).

**Fig. 4.**
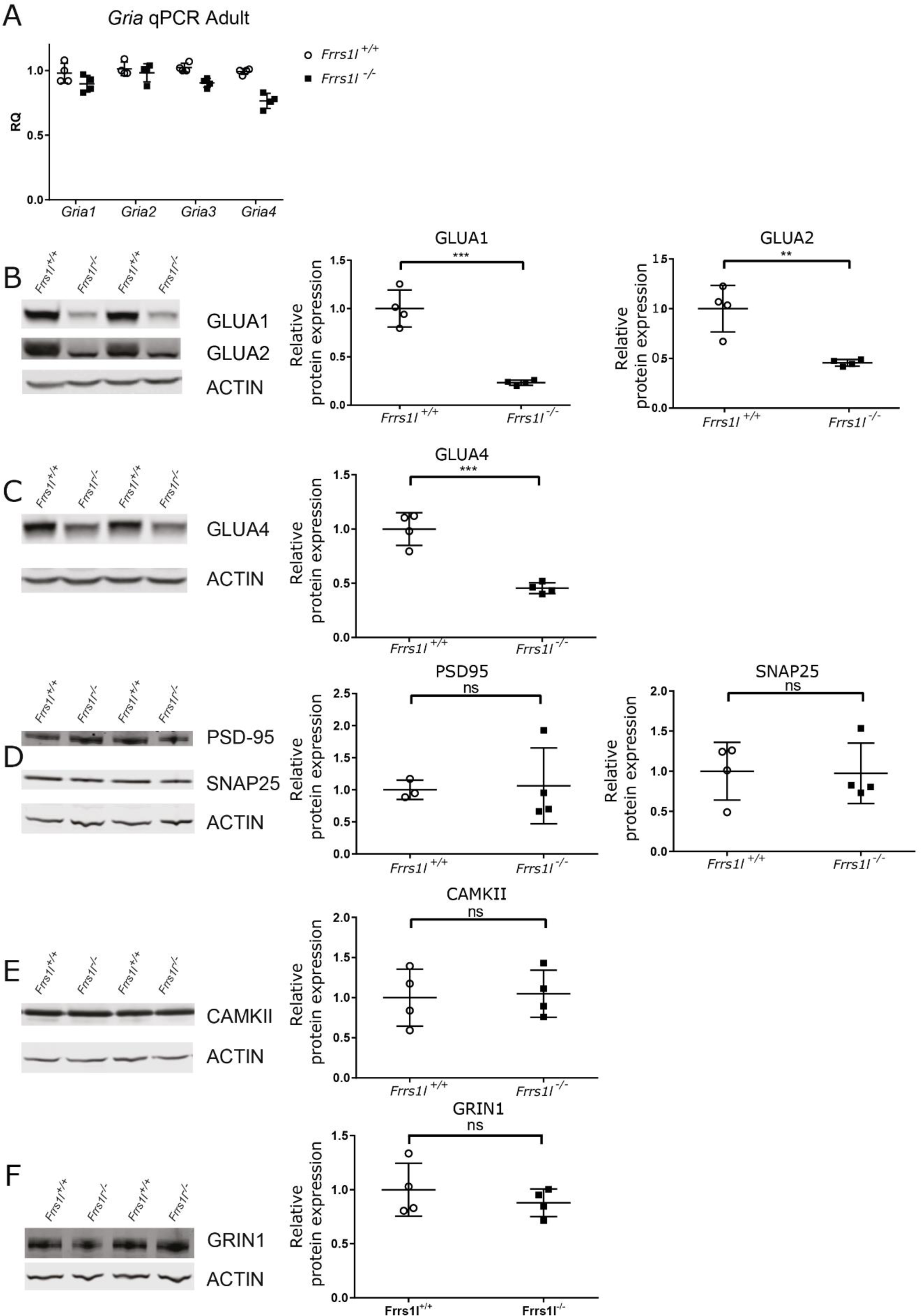
Frrs1l deficiency leads to changes in AMPA receptor subunit levels with no change in gene expression. qPCR for Gria1-4 show no changes between Frrs1l^-/-^ and wild type controls for Gria1 and Gria2, with only slight but significant changes for Gria3 and Gria4 (p<0.05 and p<0.001 respectively) (A). GLUA1, GLUA2 (B) and GLUA4 (C) immunoreactivity are all significantly lower in Frrs1l^-/-^ adult cerebellum (P<0.001, P<0.01 and P<0.001 respectively). The pattern of GLUA2 and GLUA4 differ between Frrs1l^-/-^ and wild-type controls, with a diffuse band present for GLUA2 in wild-types but only a thin band in Frrs1l^-/-^, GLUA4 blots also indicate changes in immunoreactive band mobility. Other synaptic proteins, PSD-95, SNAP25 (D) and CAMKII (E) and NMDAR (GRIN1) (F), remain unchanged between Frrs1l^-/-^ and wild-type controls. Data analysed using t-test (Frrs1l^+/+^; n=4, Frrs1l^-/-^ n=4).

To confirm that there was a specific loss of AMPA receptors in homozygotes rather than a general loss in synaptic number, we compared the levels of several standard synaptic proteins in adult cerebellum and P0 brain. Interestingly, we found no differences in expression of any of the synaptic proteins assessed (CAMKII, SNAP25, PSD95 and NMDA receptor) at any time point between wild-type and *Frrs1l*^-/-^ brains (Fig. 4D-F and S3C and D). To investigate further, we counted the proportion excitatory synapses in the hippocampus and found no differences in synapse number between wild-type and *Frrs1l*^-/-^ brains (Fig. S4). Collectively, these data show that there is a significant reduction in AMPA receptor subunit levels present in adult *Frrs1l*^-/-^ mice while synaptic numbers and levels of several key synaptic markers remain unchanged. Similar changes in protein level are observed in *Frrs1l*^-/-^ P0 brain, which points to a developmental defect rather than a progressive degenerative change.

### AMPA receptor glycosylation is incomplete leading to cytoplasmic retention in Frrs1l^-/-^ mice

Given that AMPA receptor protein levels were reduced while their transcriptional activity was unaffected, we concluded that these deficits in *Frrs1l*^-/-^ brain are most likely due to specific alterations in translational or posttranslational mechanisms. Interestingly, we observed that GLUA2 and GLUA4 band mobilities in western blots differ in wild-type and *Frrs1l*^-/-^ mice, with a more diffuse band in GLUA2 wild-type compared to in *Frrs1l*^-/-^ and a faster-running band seen in GLUA4. These differences could be related to differences in post-translational modification caused by a deficiency in this process in *Frrs1l*^-/-^ mice. As part of the maturation process of the AMPA receptor complex, the glycosylation of AMPA receptors goes through a series of modifications. Initially N-linked high mannose glycans are added to GLUA2 and GLUA4 in the endoplasmic reticulum (ER). Subsequently, AMPA receptor complexes are transported to the Golgi apparatus where glycans are clipped and modified to create more complex N-linked glycans. Assessing glycosylation state allows us not only to determine the extent of AMPA receptor glycosylation but also to establish the subcellular localisation of the AMPA receptor complex (Tucholski *et al.*, 2013, 2014).

We assessed the glycosylation status of GLUA2 and GLUA4 receptor subunits in control and mutant samples. Lysates from adult cerebellum were digested with endoglycosidase H (ENDO-H) and peptide:N-glycosidase F (PNGase) enzymes which cleave either immature glycosylated moieties, such as high mannose glycans, or all glycosylated forms, respectively (Fig. 5A). We found that, when digested with PNGase, all immunoreactive bands showed an apparent size shift, indicating that the subunits are typically glycosylated in both wild-type and *Frrs1l*^-/-^ brain. When incubated with ENDO-H, in wild-type mice, only a small proportion of the GLUA2 and GLUA4 were digested, amounting to approximately 18% of total protein for both. Therefore the majority of GLUA2 and GLUA4 in wild-type is insensitive to ENDO-H and thus must be maturely glycosylated. Conversely, in *Frrs1l*^-/-^ a greater proportion (*p<0.01*) of the GLUA2 and GLUA4 was digested with ENDO-H, amounting to approximately 65% of GLUA2 and 45% of GLUA4 total protein. This demonstrates a higher level of immature glycosylation of the receptor subunits in the absence of FRRS1L. Based on previous work following AMPA receptor localisation and glycosylation (Tucholski *et al.*, 2013, 2014), this result also suggests that GLUA2 and GLUA4 AMPA receptor processing is stalled at the level of the Golgi apparatus in *Frrs1l*^-/-^ mice.

**Fig. 5.**
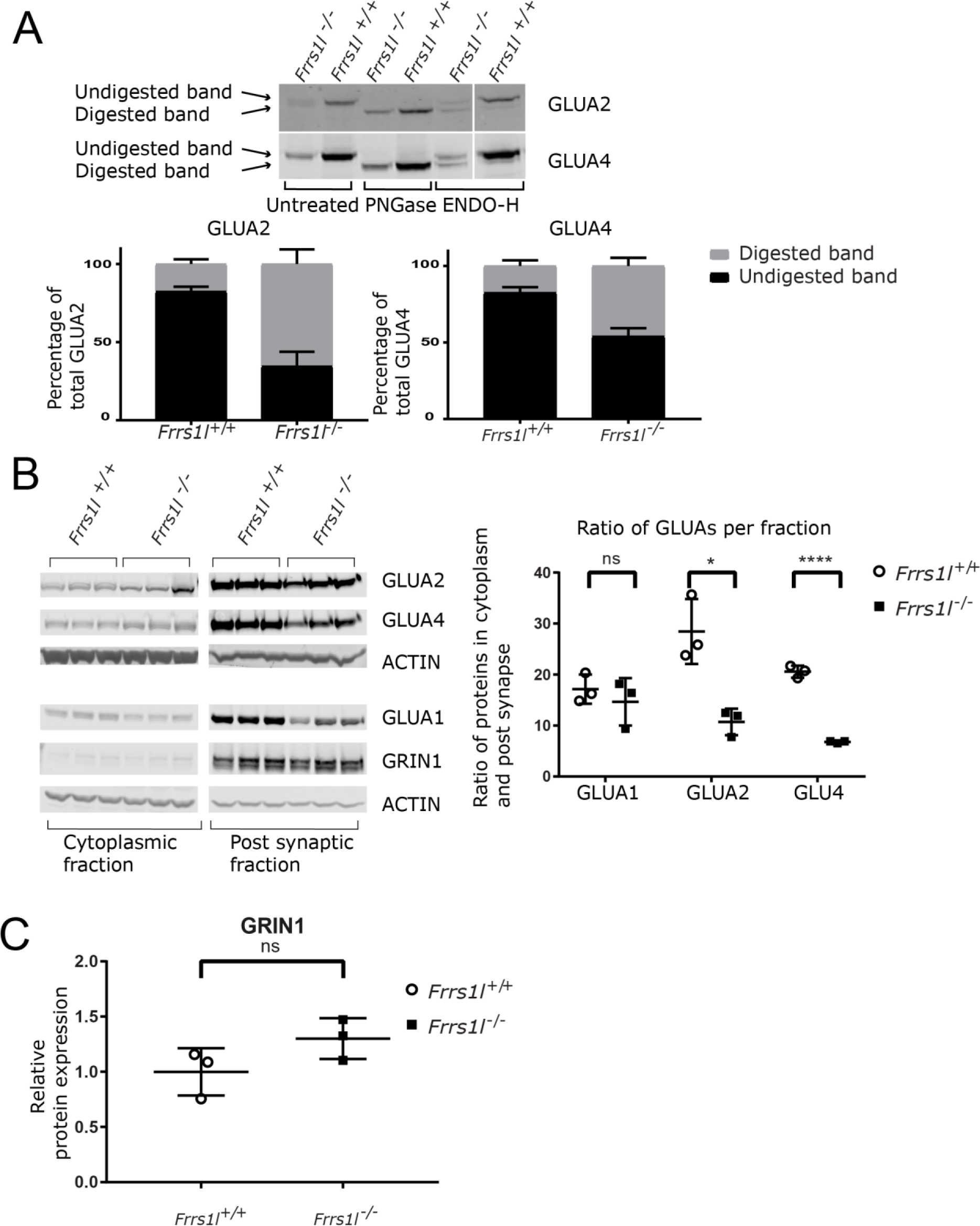
AMPA receptors have altered glycosylation state and are mislocalised in the cytoplasm in Frrs1 mice. Posttranslational glycosylation state is altered in Frrs1l^-/-^, demonstrated by differential digestion with glycosylation sensitive enzymes when compared to wild-type littermates. Frrs1l^-/-^ show greater sensitivity of GLUA2 and GLUA4 digestion with ENDO-H (p<0.01), approximately 18% of wild-type GLUA2 and GLUA4 is digested compared to approximately 65% and 45% respectively in Frrs1l^-/-^. Localisation of AMPA receptors is altered, with increased amount of AMPA receptor in the cytoplasm rather than the synapse in Frrs1l^-/-^. GLUA1 proportions in the cytoplasm and the synapse were not altered, and overall levels of GRIN1 were unchanged between Frrs1l^-/-^ and wildtype controls. Data analysed using students t-test (n=3 Frrs1l^-/-^, n=3 Frrs1l^+/+^). Percentage of total GLUA2 and 4 (A) are mean and standard deviation.

This increase in immaturely glycosylated AMPA receptors in *Frrs1l*^-/-^ mice might result in a reduction in receptor levels at the synaptic membrane. To investigate, we carried out synaptic fractionation to determine the proportional levels of AMPA receptor in the cytoplasmic and post-synaptic membrane fractions of adult forebrain. These data show that more than double the amount of both GLUA2 and GLUA4 are retained in the cytoplasm of *Frrs1l*^-/-^ compared to wild-type, leading to a reduction in the post-synaptic fraction (Fig. 5B). Interestingly, for GLUA1, levels were proportionally lower in the post-synaptic fraction in mutant brain without evidence for retention in the cytoplasmic fraction. Additionally, NMDAR levels in the post-synaptic fraction were not affected by *Frrs1l* deficiency (Fig 5C).

In conclusion, these results reveal that the loss of FRRS1L leads to a specific reduction in the levels of the AMPA receptor subunit proteins at the synapse *in vivo* without obvious changes in other synaptic components. Functional FRRS1L is necessary for the mature glycosylation of at least GLUA2 and GLUA4. Loss of FRRS1L leads to incomplete post-translational processing of AMPA receptors, increased retention of AMPA receptors in the cytoplasmic fraction and, consequently, a decrease in levels of functional AMPA receptors at the synapse.

## Discussion

In previous work we described several families with homozygous mutations in *FRRS1L* (Madeo *et al.*, 2016). The main symptoms in the affected children are encephalopathy, epilepsy and progressive choreoathetosis. All children have severe intellectual disability with no expressive speech, impaired volitional movement and chorea developing into hypokinesia and seizures. This is a rare disease only recently characterized, although it is expected that there will be more cases in the near future, especially considering the gene is now included in the screening for infantile epilepsy and dyskinesia (GTR test ID GTR000551789.3).

Our data demonstrate that complete lack of *Frrs1l* has substantial effects on post-natal survival, as well as body weight, motor co-ordination, activity, sleep, effects on cognition and abnormal EEG. Anomalies are present from an early age with no progressive deterioration, suggesting a neurodevelopmental defect, rather than an age-associated neurodegenerative disorder. Interestingly *Frrs1l*^-/-^ mice show sleep disturbances. Although sleep disturbances have not been documented in human patients with mutations in *FRSS1L*, it is well known that sleep disruption is often a feature of numerous neurological disorders, particularly epileptic disorders (Crespel, Baldy-Moulinier and Coubes, 1998; Kothare and Kaleyias, 2010). Importantly, we report the occurrence of behavioural seizures in these mice, although we were unable to capture them on EEG recording, which instead showed prominent runs of delta waves. Thus, all these phenotypic features resemble the symptoms seen in patients carrying homozygous mutations in the *FRRS1L* gene, making these mice a very useful model of disease. The use of these mice allowed us to provide compelling *in vivo* evidence that FRRS1L is critical for AMPA receptor complex maturation, defects in which result in dramatic phenotypic effects in mice.

In *Frrs1l*^-/-^ mice we found a highly significant reduction in the levels of core AMPA receptor proteins from birth through to adulthood, and importantly, AMPA receptor subunits lack complete posttranslational processing and mature glycosylation. Thus, these data provide *in vivo* evidence that FRRS1L is crucial for the correct biogenesis and maturation of AMPA receptors, which elicits dramatic motor alterations, dyskinesia phenotypes, and abnormal electrographical activity in *Frrs1l*^-/-^ mice.

Previous work using a CRISPR/Cas9 deletion of *Frrs1l* in mouse primary neurons shows that it leads to an overall reduction in GLUA1 levels (Han *et al.*, 2017). Here we confirm that GLUA1 levels are lower *in vivo* but we extend this observation to other GLUAs. Crucially, we show that the low AMPA receptor levels are not a consequence of degeneration nor are they associated with a general reduction in synaptic number in *Frrs1l*^-/-^ mice. Thus the role of FRSS1L seems to be specific in the maturation of AMPA receptor complexes. Importantly we found that it is not only the levels of AMPA receptors that are low, but also their maturation (glycosylation) and location at synapses. AMPA receptors which are not fully glycosylated are not functional (Tomita *et al.*, 2003; Tucholski *et al.*, 2014), and so the presence of functional receptor complexes in the membrane are substantially deficient in mutants. We also provide evidence that FRSS1L has a critical role in the glycosylation/maturation of GLUA2 and GLUA4. It is interesting to note that the glycosylation of GLUA2 and GLUA4 is not totally abolished, indicating that there could be partial compensation for FRRS1L by another protein or perhaps a parallel mechanism, independent of FRRS1L, which may be involved in AMPA receptor biogenesis.

Supporting these data, the *Frrs1l*^-/-^ mouse phenotype resembles that of null mutations in other genes associated with synthesis, transport and stability of AMPA receptors, such as *Cpt1c* (Carrasco *et al.*, 2013), *Shank3* (Wang *et al.*, 2011) and *Stargazin* (Letts *et al.*, 1998; Menuz and Nicoll, 2008). CPT1C is proposed to play a similar role to FRRS1L in AMPA receptor biogenesis (Brechet *et al.*, 2017). As might be expected, many phenotypic similarities can be found in knockouts of *Cpt1c* and *Frrs1l*. Carrasco et al (Carrasco *et al.*, 2013) demonstrate that *Cpt1c* knockout mice have poor co-ordination, reduced latency to fall from a rotarod, ataxia and reduced grip strength. Conversely, the *Cpt1c* knockout mice show hypoactivity, whereas we demonstrate that *Frrs1l*^-/-^ mice are hyperactive. It is possible that this inconsistency is due to the different methods of activity test measurement in the two studies. Interestingly, *Cpt1c* knockout mice show reduced levels of AMPA receptor subunit proteins, with no change in AMPA receptor gene expression by qPCR, mirroring the results seen in *Frrs1l*^-/-^ mice (Fadó *et al.*, 2015) and substantiating the argument that these two proteins are involved in the same process. Null mutations in mouse *Shank3*, a scaffold protein in the post synaptic density, results in abnormal foot placement and reduced latency to fall from a rotarod, both of which are seen in *Frrs1l*^-/-^ mice. However *Shank3* mice also show decreased locomotion which is contrary to the phenotype of *Frrs1l*^-/-^. The Stargazin mouse has a mutation in PSD-95, a member of the AMPA receptor complex, which causes a general decrease in AMPA receptor function and phenotypes that overlap with *Frrs1l* mutants, specifically ataxia and impaired co-ordination (Letts *et al.*, 1998; Menuz and Nicoll, 2008). Stargazin mice also have a similar change in glycosylation of GLUA2 as seen in *Frrs1l*^-/-^. It is noticeable that one of the main features of patients with homozygous mutations in *FRRS1L* and in the stargazin mouse model is the presence of seizures. Crucially, we are able to detect behavioural seizures in this study of *Frrs1l*^-/-^ mice.

In summary, we provide evidence for the validity of the *Frrs1l*^-/-^ mouse as a model of disease, expressing phenotypic features that resemble many of the clinical symptoms in patients. At the molecular level we have demonstrated that *FRSS1L* has a fundamental role in AMPA receptor biology, impacting in the total AMPA receptor levels as well as a reduction in the proportion of AMPA receptors available at synapses. This mouse is potentially an important pre-clinical model not only to support the development of therapeutics for such patients, but also as a valuable resource to further understand the complexities of AMPA receptors and glutamate signalling in the brain.

## Materials and Methods

### Mice

All mice *(Mus musculus*) were maintained and studied in accordance with UK Home Office legislation and local ethical guidelines issued by the Medical Research Council (Responsibility in the Use of Animals for Medical Research, July 1993; home office licenses 30/2890 and 30/3384). Mice were fed *ad libitum* on a commercial diet (SDS Rat and Mouse No. 3 Breeding diet, RM3) and had free access to water (9–13 ppm chlorine). Mice were kept under controlled light (light 7am–7pm, dark 7pm–7am), temperature (21±2°C) and humidity (55±10%) conditions.

*Frrs1l^tm1a/+^* mice were derived from C57BL6/NTac ES cells (Skarnes et al, 2011). The null allele (tm1b) was created by carrying out an IVF using *Frrs1l^tm1a/+^* sperm and C57BL6/NTac oocytes. Soluble cell permeable cre (TAT-Cre (Tat-NLS-Cre, HTNC, HTNCre), Excellegen, Rockville, USA) was added to two cell *Frrs1l^tm1a/+^* embryos to generate the *Frrs1l^tm1b/+^*allele. The cre excises the selection cassette and exon 3 of the *Frrs1l* gene, creating a null allele (https://www.i-dcc.org/imits/targ_rep/alleles/14093/allele-image?simple=true.jpg). Following washing to remove the soluble cre, the IVF procedure was completed as normal. The *Frrs1l^tm1b/+^* were crossed to C57BL6/NTac and then intercrossed to create *Frrs1l^tm1b/tm1b^*, *Frrs1l^tm1b/+^* and *Frrs1l^+/+^* cohorts. Hereafter *Frrs1l^tm1b/tm1b^* will be referred to as *Frrs1l*^-/-^.

### Behavioural phenotyping tests

For all behavioural phenotyping tests, mice were taken to the test room at least 20 minutes prior to the start of the test to acclimatise. Phenotyping equipment was cleaned with 70% ethanol/IMS or 2% Distel between tests.

Investigators were blind to genotype during all phenotyping tests.

All phenotyping except fear conditioning was carried out on female mice due to issues of reduced viability. SHIRPA, grip strength and home cage activity monitoring were assessed at 3, 6 and 9 months. All other tests were carried out on only one occasion at the ages indicated below. For behavioural tests, sample size was calculated using power equations based on previous data obtained on C57BL/6NTac mice. Sample sizes in the later time points are smaller due to the loss of several mice with welfare concerns.

#### Fear conditioning

Fear conditioning was carried out on a separate cohort of male mice prior to the rest of the study (n=7 C57BL/6NTac; n=8 *Frrs1l*^+/−^, n=7 *Frrs1l*^-/-^). These mice were part of the IMPC phenotyping pipeline, the remainder of the IMPC phenotyping data is published on the IMPC data portal (IMPC, http://www.mousephenotype.org/, Koscielny *et al.*, 2014).

#### Open Field Activity

Open field activity was used to assess locomotion in a novel environment. At 10 weeks of age (+/-1 week), mice were placed in square arenas (44×44cm) in a small testing room. A minimum of two and a maximum of four mice were tested at one time, one mouse per arena. Lighting was set at 150-200 lux. Mice were video tracked for 20 minutes and data analysed using Ethovision XT software (Noldus, Netherlands) and parameters such as distance moved, velocity and duration moving were recorded in various zones over the entire 20 minute period (Joyce *et al.*, 2016). (n=7 *Frrs1l* ^+/+^; n=10 *Frrs1l* ^-/-^).

#### SHIRPA

A semi-quantitative assessment was carried out using a modified SmithKline Beecham, Harwell Imperial College, Royal London Hospital phenotype assessment (SHIRPA) protocol. Behaviour and dysmorphology parameters were recorded as previously described (Masuya *et al.*, 2005) (n=8 *Frrs1l* ^+/+^; n=9 *Frrs1l* ^-/-^ at three months, n=8 *Frrs1l* ^+/+^; n=9 *Frrs1l*^-/-^ at 6 months, n=8 *Frrs1l* ^+/+^; n=7 *Frrs1l* ^-/-^at 9 months).

#### Grip Strength

Grip strength was assessed at 13 weeks of age (+/-1 week) using the Grip Strength Test (BioSeb, Chaville, France). Readings were taken from all four paws, three times per mouse at each age, as per manufacturer’s instructions (Joyce *et al.*, 2016). (n=8 *Frrs1l* ^+/+^; n=8 *Frrs1l* ^-/-^ at three months, n=8 *Frrs1l* ^+/+^; n=8 *Frrs1l* ^-/-^ at 6 months, n=6 *Frrs1l*^-/-^ n=6 *Frrs1l*^+/+^ at 9 months).

#### Home cage analysis

Group housed animals were monitored as described (Bains *et al.*, 2016). Briefly, group housed mice were tagged with RFID micochips at 9 weeks of age and placed in the Home Cage Analysis system (Actual Analytics, Edinburgh) which captured mouse behaviour using both video tracking and location-tracking using RFID co-ordinates (Bains *et al.*, 2016). (n=8 *Frrs1l* ^+/+^; n=8 *Frrs1l* ^-/-^ at all time points).

#### Locotronic

Paw placement was analysed using Locotronic (Intelli-Bio, France) at 13 weeks of age (+/-1 week). Briefly, animals were assessed as they moved down a corridor with a horizontal ladder as its base. Animals were motivated to travel from a lighter starting area at one end to a darker finish area at the other end (bars: 3 mm diameter; spaced by 7 mm). Infrared sensors above and below each bar space recorded any errors of paw placement. Trials were discounted if the mouse took more than 30s to move to the finish after exiting the start area (n=8 *Frrs1l* ^+/+^; n=7 *Frrs1l* ^-/-^).

#### Rotarod

To assess co-ordination and motor learning, 22-week-old mice (+/-2 weeks) were placed on an accelerating rotarod (Ugo Basile), with rotor speed increasing from 4rpm up to 40rpm over a five-minute period. The time taken for the mouse to fall from the rod was recorded. This was repeated three times on one day with a 15 minute inter trial interval (Corrochano *et al.*, 2012) (n=8 *Frrs1l* ^+/+^; n=9 *Frrs1l* ^-/-^).

#### Y-maze

A forced alternation y-maze test was used to evaluate short term working memory in mice at 14-20 weeks of age. Mice were placed in a y-maze with access to one arm blocked, they were then free to explore the start arm and the ‘familiar’ arm for ten minutes. Mice were returned to the home cage for a two minute inter trial interval, during which the maze was cleaned to remove odour cues. The mice were then returned to the maze, with access to all three arms open for five minutes. Mice were video tracked at all times using Ethovision software (Noldus, Netherlands) (Sanderson and Bannerman, 2012). (n=7 *Frrs1l* ^+/+^; n=7 *Frrs1l* ^-/-^).

#### Motor function assessment by wheel running

For further assessment of motor function, 45-50 week old mice were singly housed and placed in cages containing a running wheel as previously described (Mandillo *et al.*, 2014) (TSE systems Bad Homburg, Germany). Number of rotations, time running, number of bouts and speed were measured. After two weeks with the standard wheel, this was replaced with a complex wheel which had specific rungs removed in order to test co-ordination and learning. Parameters were recorded for a further week (n=5 *Frrs1l* ^+/+^; n=5 *Frrs1l* ^-/-^).

### Passive Infrared screen for immobility defined sleep (PIR)

At 1 year, mice were analysed for circadian activity and immobility defined sleep using the COMPASS system as described (Brown *et al.*, 2016). Mice were individually housed and data captured for 5 days in a 12:12 LD cycle, followed by 9 days in constant darkness. Data analysis was performed using custom python scripts and excels sheets, developed in house. Circadian analysis was performed by converting activity data from PIR to AWD files for analysis on Clocklab (Actimetrics, Illinois) or Actiwatch Sleep analysis software (CamNtech, Cambridge) (n=5 *Frrs1l* ^+/+^; n=6 *Frrs1l* ^-/-^).

### Quantitative PCR analysis

RNA extraction from P0 brain tissue or cerebellum of 14 month old mice was performed using an RNeasy kit (Qiagen) (n=5 *Frrs1l* ^+/+^; n=5 *Frrs1l* ^-/-^ at P0, n=4 *Frrs1l* ^+/+^; n=4 *Frrs1l* ^-/-^ at 14 months). cDNA synthesis was performed using the High Capacity cDNA RT kit (ThermoFisher Scientific) starting with 2µg of total RNA. cDNA for qPCR amplification was used at a final concentration of 10 ng per well. All the reactions were run in triplicate. Fast Sybr Green mastermix from ThermoFisher Scientific was used and the reactions had a final volume of 20µl. Primers were at a final concentration of 360nM. Primers were designed to span exon-exon boundaries and are listed in Supplemental Table 1. Fold changes were calculated using the 2-ddCt method using the 7500 Software v2.0.6 and normalized using S16 endogenous reference genes relative to WT genotype (Livak and Schmittgen, 2001).

### Immunoblot analysis

Adult cerebellum or P0 whole brains were bisected and one fraction homogenized in RIPA buffer (150 mM NaCl, 1% NP40, 0.5% Na deoxycholate, 0.1% SDS, 50 mM Tris pH 7.5) with phosphatase and protease inhibitor cocktails (Roche), using lysing matrix tubes D (MP Biomedicals, Germany) and a Fast-Prep-24 homogenizer at 4°C. Homogenates were centrifuged at 12,000 *g* 4°C for 20 min. 30 μg of soluble fractions were resolved by SDS– PAGE (NUPAGE system, Invitrogen) and transferred to Nitrocelulose membranes (Millipore) for western-blot analysis. The following primary antibodies were used: rabbit monoclonal anti-b-actin (1/3000 Sigma A2066); mouse anti-α tubulin (1/3000 Sigma T9026), rabbit anti-GLUA1 (1/1000 Millipore 1504) mouse anti-GLUA2 (1/800 Millipore MAB397), rabbit anti-GLUA4 (1/1000 Millipore AB1508), mouse anti-CAMKII (1/200 Proteintech 20665-1) mouse anti-SNAP25 (1/500 Biolegend 836304), rabbit anti-PSD-95 (1/1000 Cell Signalling 3450T), and mouse anti-GRIN1 (for NMDA receptor, 1/1000 Novus, NB300-118).

Protein was visualized using anti-mouse (P/N 926-68070) or anti-rabbit (P/N 926-32211) secondary antibodies IRDye^®^ (Li-Cor Biosciences) at 1:10000 dilutions and quantified using the scanning infrared Odyssey imaging system CX (Li-Cor Biosciences).

All antibodies used are previously validiated, published and purchased from commercial suppliers.

### Glycosylation assay

Bisected adult cerebellum was homogenised in RIPA buffer as above. 60ug of soluble fraction was de-natured with glycoprotein denaturing buffer for 10 minutes at 100 degrees, followed by immediate immersion in ice. Samples were divided into three aliquots and glycobuffer added, followed by Endo-H (QABio, E-EH02), PNGase (New England Biolabes, P0704S) or water. All samples were incubated at 37 degrees for 1 hour. Half of each sample, containing 10μg of protein, was resolved by SDS–PAGE (NUPAGE system, Invitrogen) and transferred to Nitrocellulose membranes for Western blot analysis.

### Post-Synaptic Fractionation enrichment

Synaptic fractionation was conducted following a modified protocol using a single cerebral hemisphere homogenized using a Dounce homogenizer in Syn-PER™ Synaptic Protein Extraction Reagent (Thermo Scientific, IL, USA) with phosphatase and protease inhibitor cocktails (Roche). Homogenates were cleared by centrifugation at 1200 x g for 10 min and then at 15000 x g for 20 min at 4 degrees. The supernatant contains the cytosolic fraction and the pellet the crude synaptosomal fraction. The pellet was resuspended in syn-PER lysis buffer with 0.1mM CaCl2 and 2% Triton X-100, 40mM Tris, pH6 and incubated on ice, with gentle agitation for 30 minutes. Following a centrifugation at 40,000 x g for 30 minutes at 4°C, the pellet was washed with 1% Triton X-100, 20mM Tris, pH6. Then the sample was centrifuged again, resuspended in 1% Triton x-100, 20mM Tris, pH8 and incubated on ice with gentle agitation for 30 minutes. After another centrifugation at 40,000 x g for 30 minutes at 4°C the pellet, containing the post-synaptic density, was resuspended in 1% Triton x-100, 20mM Tris, pH8, precipitated by adding 10 volumes of ice cold acetone at °20 C overnight and subsequently centrifuged at 15,000 x g for 30 minutes, at 4°C. The pellet containing the post-synaptic fraction was resuspended in 5 % SDS. After protein quantification with a DC assay (BioRad), 15 μg of protein from the post-synaptic fraction and 20 μg from the cytoplasmic fraction were resolved in precast 3-8% SDS-PAGE gels and transferred to Nitrocellulose membrane (Invitrogen) for Western blot analysis.

### Synapse Counts

Formalin fixed wax embedded sections of whole brain were dewaxed, the antigen was unmasked using sodium citrate buffer solution (ph 6.0) at 80 °C for 30 minutes. Sections were next washed in phosphate-buffered saline (PBS, Ph 7.4) and processed for immunofluorescence. After a blocking step in PBS containing 0.05% Triton X-100 and 10% normal goat serum (NGS), sections were incubated overnight at 4 °C with the antibodies anti-VGLUT1 (1:200, Synaptic System. Antibody was diluted in PBS with 3% normal goat serum and 0.05% Triton X-100. Sections were then washed in PBS (4 x 10 min) and incubated for 1 h at room temperature with a secondary anti-rabbit antibody conjugated with Alexa Fluor 488 (Invitrogen). After several PBS rinses, sections were mounted on glass slides and observed with a Zeiss LSM 700 confocal microscope (Carl Zeiss AG). Confocal z-stacks covering the whole depth of the slices (1024 x 1024 pixels) spaced by 1.05 µm were acquired at 63 x. VGLUT1 positive puncta were analysed on confocal images using Fiji software. Caudal sections were used to analyse both stratum oriens and radiatum of the CA1 region of the hippocampus.

### Telemetry EEG

Electroencephalography (EEG) measurements were conducted in male mice, around 30g in weight, and age between 4 and 6 months (n=3 *Frrs1l^-/-^,* n=2 *Frrs1l^/+/+^*, and n=2 C57BL/6NTac). Wireless EEG transmitters (A3028A, Open Source Instruments, USA) were implanted subcutaneously with a subdural intracranial recording electrode positioned above the right frontal lobe (0.5mm anterior to Bregma and 0.5mm medial-lateral). The second electrode was implanted in between the frontal and occipital lobes, above the right motor cortex (1 – 1.5mm posterior to Bregma and 3 - 4 mm medial-lateral). The animals were able to freely move while assessed in their home-cage environment for 5 to 21 days. EEG was recorded after at least 3 or more post-surgery recovery days to ensure no residual anaesthesia effect. Dataquest software (Neuroachieve v. 8.5.20, Open Source Instruments, USA) was used to acquire and analyse the EEG data. EEG activity was sampled at 512 Hz at between 0.3Hz and 160Hz. Representative high amplitude signals were screened visually and fast fourier transform was used to determine EEG amplitude and frequency.

### Statistical analysis

Estimates of Mendelian inheritance of all genotypes were assessed using a chi-squared test. Analysis of categorical data from SHIRPA was completed using Fishers exact test. Grip strength test, rotarod, home cage analysis, time spent asleep, sleep bouts over time, sleep fragmentation and motor function wheel running data were assessed using repeated measures analysis of variance (ANOVA) with Sidak post-hoc analysis. Body weight was assessed using two-way ANOVA with post hoc comparisons. Forced alternation y-maze data were assessed using the Student’s t-test. Open field data were analysed using the Student’s t-test with Welch’s correction for unequal variance. Graphpad Prism and R software were used for statistical analysis.

All graphs show absolute values with mean and standard deviation, unless specified.

## Acknowledgements

The authors thank MLC animal technicians and genotyping for technical support. Chris Esapa for advice on glycosylation experiments.

## Author contributions

M.S, A.A-A, S.W, P.M.N., and S.C. conceived the project, designed the experiments and wrote the manuscript. M.S, P.L, G.B., E.C, S.R., C.D., and S.C. carried out research. P.O., D.K., M.K., contributed to the writing and analysis of the experiments. All authors checked for scientific content and contributed to the final drafting of the manuscript.

## References

Bains, R. S. et al. (2016) ‘Analysis of Individual Mouse Activity in Group Housed Animals of Different Inbred Strains using a Novel Automated Home Cage Analysis System’, Frontiers in Behavioral Neuroscience, 10(June), pp. 1–12. doi: 10.3389/fnbeh.2016.00106.

Brechet, A. et al. (2017) ‘AMPA-receptor specific biogenesis complexes control synaptic transmission and intellectual ability’, Nature Communications, 8(May), p. 15910. doi: 10.1038/ncomms15910.

Brown, L. A. et al. (2016) ‘COMPASS: Continuous Open Mouse Phenotyping of Activity and Sleep Status’, Wellcome Open Res, 1(May), p. 2. doi: 10.12688/wellcomeopenres.9892.1.

Carecchio, M. and Mencacci, N. E. (2017) ‘Emerging Monogenic Complex Hyperkinetic Disorders’. Current Neurology and Neuroscience Reports.

Carrasco, P. et al. (2013) ‘Carnitine palmitoyltransferase 1C deficiency causes motor impairment and hypoactivity’, Behavioural Brain Research. Elsevier B.V., 256, pp. 291–297. doi: 10.1016/j.bbr.2013.08.004.

Chen, L. et al. (2000) ‘Stargazin regulates synaptic targeting of AMPA receptors by two distinct mechanisms.’, Nature, 408(6815), pp. 936–43. doi: 10.1038/35050030.

Corrochano, S. et al. (2012) ‘α-Synuclein levels modulate Huntington’s disease in mice’, Human Molecular Genetics, 21(3), pp. 485–494. doi: 10.1093/hmg/ddr477.

Crespel, A., Baldy-Moulinier, M. and Coubes, P. (1998) ‘The relationship between sleep and epilepsy in frontal and temporal lobe epilepsies: Practical and physiopathologic considerations’, Epilepsia, 39(2), pp. 150–157. doi: 10.1111/j.1528-1157.1998.tb01352.x.

Davies, B. et al. (2017) ‘A point mutation in the ion conduction pore of AMPA receptor GRIA3 causes dramatically perturbed sleep patterns as well as intellectual disability’, Human Molecular Genetics, 26(20), pp. 3869–3882. doi: 10.1093/hmg/ddx270.

Erlenhardt, N. et al. (2016) ‘Porcupine Controls Hippocampal AMPAR Levels, Composition, and Synaptic Transmission’, Cell Reports. The Authors, 14(4), pp. 782–794. doi: 10.1016/j.celrep.2015.12.078.

Fadó, R. et al. (2015) ‘Novel regulation of the synthesis of α-Amino-3-hydroxy-5-methyl-4-isoxazolepropionic Acid (ampa) receptor subunit glua1 by carnitine palmitoyltransferase 1C (CPT1C) in the Hippocampus’, Journal of Biological Chemistry, 290(42), pp. 25548–25560. doi: 10.1074/jbc.M115.681064.

Han, W. et al. (2017) ‘Ferric Chelate Reductase 1 Like Protein (FRRS1L) Associates with Dynein Vesicles and Regulates Glutamatergic Synaptic Transmission’, Frontiers in Molecular Neuroscience, 10(December). doi: 10.3389/fnmol.2017.00402.

Joyce, P. I. et al. (2016) ‘Deficiency of the zinc finger protein ZFP106 causes motor and sensory neurodegeneration’, Human Molecular Genetics, 25(2), pp. 291–307. doi: 10.1093/hmg/ddv471.

Kato, A. S. et al. (2010) ‘Hippocampal AMPA Receptor Gating Controlled by Both TARP and Cornichon Proteins’, Neuron, 68(6), pp. 1082–1096. doi: 10.1016/j.neuron.2010.11.026.

Koscielny, G. et al. (2014) ‘The International Mouse Phenotyping Consortium Web Portal, a unified point of access for knockout mice and related phenotyping data’, Nucleic Acids Research, 42(D1), pp. 802–809. doi: 10.1093/nar/gkt977.

Kothare, S. V. and Kaleyias, J. (2010) ‘Sleep and epilepsy in children and adolescents’, Sleep Medicine. Elsevier B.V., 11(7), pp. 674–685. doi: 10.1016/j.sleep.2010.01.012.

LaBar, K. S. and Cabeza, R. (2006) ‘Cognitive neuroscience of emotional memory’, Nature Reviews Neuroscience, 7(1), pp. 54–64. doi: 10.1038/nrn1825.

Lalonde, R. and Strazielle, C. (2011) ‘Brain regions and genes affecting limb-clasping responses’, Brain research reviews, 67, pp. 252–259.

Letts, V. A. et al. (1998) ‘The mouse stargazer gene encodes a neuronal Ca 2 + - channel γ subunit’, 19(august), pp. 340–347.

Livak, K. J. and Schmittgen, T. D. (2001) ‘Analysis of relative gene expression data using real-time quantitative PCR and the 2-ΔΔCT method’, Methods, 25(4), pp. 402–408. doi: 10.1006/meth.2001.1262.

Madeo, M. et al. (2016) ‘Loss-of-Function Mutations in FRRS1L Lead to an Epileptic-Dyskinetic Encephalopathy’, American Journal of Human Genetics. American Society of Human Genetics, 98(6), pp. 1249–1255. doi: 10.1016/j.ajhg.2016.04.008.

Mandillo, S. et al. (2014) ‘Early motor deficits in mouse disease models are reliably uncovered using an automated home-cage wheel-running system: a cross-laboratory validation.’, Disease models & mechanisms, 7(3), pp. 397–407. doi: 10.1242/dmm.013946.

Masuya, H. et al. (2005) ‘Implementation of the modified-SHIRPA protocol for screening of dominant phenotypes in a large-scale ENU mutagenesis program’, Mammalian Genome, 16(11), pp. 829–837. doi: 10.1007/s00335-005-2430-8.

Menuz, K. and Nicoll, R. A. (2008) ‘Loss of Inhibitory Neuron AMPA Receptors Contributes to Ataxia and Epilepsy in Stargazer Mice’, Journal of Neuroscience, 28(42), pp. 10599–10603. doi: 10.1523/JNEUROSCI.2732-08.2008.

Sanderson, D. J. and Bannerman, D. M. (2012) ‘The Role of Habituation in Hippocampus-Dependent Spatial Working Memory Tasks : Evidence From GluA1 AMPA Receptor Subunit Knockout Mice’, 994, pp. 981–994. doi: 10.1002/hipo.20896.

Schwenk, J. et al. (2012) ‘High-Resolution Proteomics Unravel Architecture and Molecular Diversity of Native AMPA Receptor Complexes’, Neuron, 74(4), pp. 621–633. doi: 10.1016/j.neuron.2012.03.034.

Schwenk, J. et al. (2014) ‘Regional diversity and developmental dynamics of the AMPA-receptor proteome in the mammalian brain’, Neuron. Elsevier Inc., 84(1), pp. 41–54. doi: 10.1016/j.neuron.2014.08.044.

Shaheen, R. et al. (2016) ‘Epileptic encephalopathy with continuous spike-and-wave during sleep maps to a homozygous truncating mutation in AMPA receptor component FRRS1L’, Clinical Genetics, 90(3), pp. 282–283. doi: 10.1111/cge.12796.

Skarnes WC, et al (2011). ‘A conditional knockout resource for the genome-wide study of mouse gene function’. Nature. 474, pp337–342.

Tomita, S. et al. (2003) ‘Functional studies and distribution define a family of transmembrane AMPA receptor regulatory proteins’, Journal of Cell Biology, 161(4), pp. 805–816. doi: 10.1083/jcb.200212116.

Tucholski, J. et al. (2013) ‘Abnormal N-linked glycosylation of cortical AMPA receptor subunits in schizophrenia’, Schizophrenia Research, 146(1–3), pp. 177–183. doi: 10.1016/j.schres.2013.01.031.

Tucholski, J. et al. (2014) ‘Evolutionarily conserved pattern of AMPA receptor subunit glycosylation in mammalian frontal cortex’, PLoS ONE, 9(4). doi: 10.1371/journal.pone.0094255.

Wang, X. et al. (2011) ‘Synaptic dysfunction and abnormal behaviors in mice lacking major isoforms of Shank3’, Human Molecular Genetics, 20(15), pp. 3093–3108. doi: 10.1093/hmg/ddr212.

Zeisel, A et al. Molecular architecture of the mouse nervous system bioRxiv 294918; doi: https://doi.org/10.1101/294918

